# Characterization Of Yellow Root Cassava And Food Products: Investigation Of Cyanogenic Glycosides And Pro-Vitamin A

**DOI:** 10.1101/2020.04.03.024224

**Authors:** Chiemela S. Odoemelam, Benita Percival, Zeeshan Ahmad, Ming-Wei Chang, Dawn Scholey, Emily Burton, Polycarp N. Okafor, Philippe B. Wilson

## Abstract

**Objective:** Cyanide is a highly toxic compound, and the consumption of products containing cyanide is of singificant public health concern. In contrast, *β*-carotene possesses essential nutritional attributes related to human health, therefore the characterisation and quanfication of both compounds in food products is both fundamental and necessary. This investigation sought to identify the cyanide and β-carotene levels in two flours produced from the roots of two varieties of cassava (*Manihot esculenta crantz*), namely UMUCASS-38 (TMS 01/1371) and NR 8082, and their associated food products.

**Results:** The fresh tuber, raw flour and food products were analysed for levels of residual cyanide and *β*-carotene using standard analytical methods. The cyanide content of NR 8082 (18.01±0.01 ppm) and UMUCASS 38 (17.02±0.02 ppm) flours were significantly higher (*p* < 0.05) than the residual cyanide levels determined in the cookies (10.00±0.00 ppm) and cake (7.10±0.14 ppm). The levels of β-carotene determined in the sample varied significantly (*p* < 0.05). The highest levels of β-carotene (6.53±0.02 µg/g) were determined in raw roots of UMUCASS 38 while NR 8082 levels of β-carotene were 1.12±0.02 µg/g. Processing the roots into flour reduced the β-carotene content to 4.78±0.01 µg/g and 0.76±0.02 µg/g in UMUCASS 38 and NR8082 flours, respectively. Cookies and cake produced from flour derived from the UMUCASS 38 variety had 2.15±0.01 µg/g and 2.84±0.04 µg/g of β-carotene, respectively.

## Introduction

Many stems (yams and sweet potatoes) and root tubers (cassava) serve as food for humans and animals. Cassava is among the staple foods in many parts of Africa, Asia and Latin America, with its roots being one of the main sources of carbohydrates in the region. Cassava is recognised as a crop which requires low agrochemical input, as well as being one of the most draught complaisant crops. Hence, it thrives even in mediocre soils [1]. There has been a substantial increase in world production of cassava since 2001, with the peak reaching 293.01 million tons in 2015 (Fig. 1a) [2]. According to FAOSTAT [2], world cassava production for the year 2018 is estimated to be approximately 277.81 million tons. In the last 10 years, the top five countries for cassava production were Nigeria, Thailand, Democratic Republic of Congo (DRC), Brazil and Indonesia with an average of 50.98, 28.66, 26.81, 22.67 and 21.85 million tons of production respectively (Fig. 1b) [2].

**Fig. 1.**
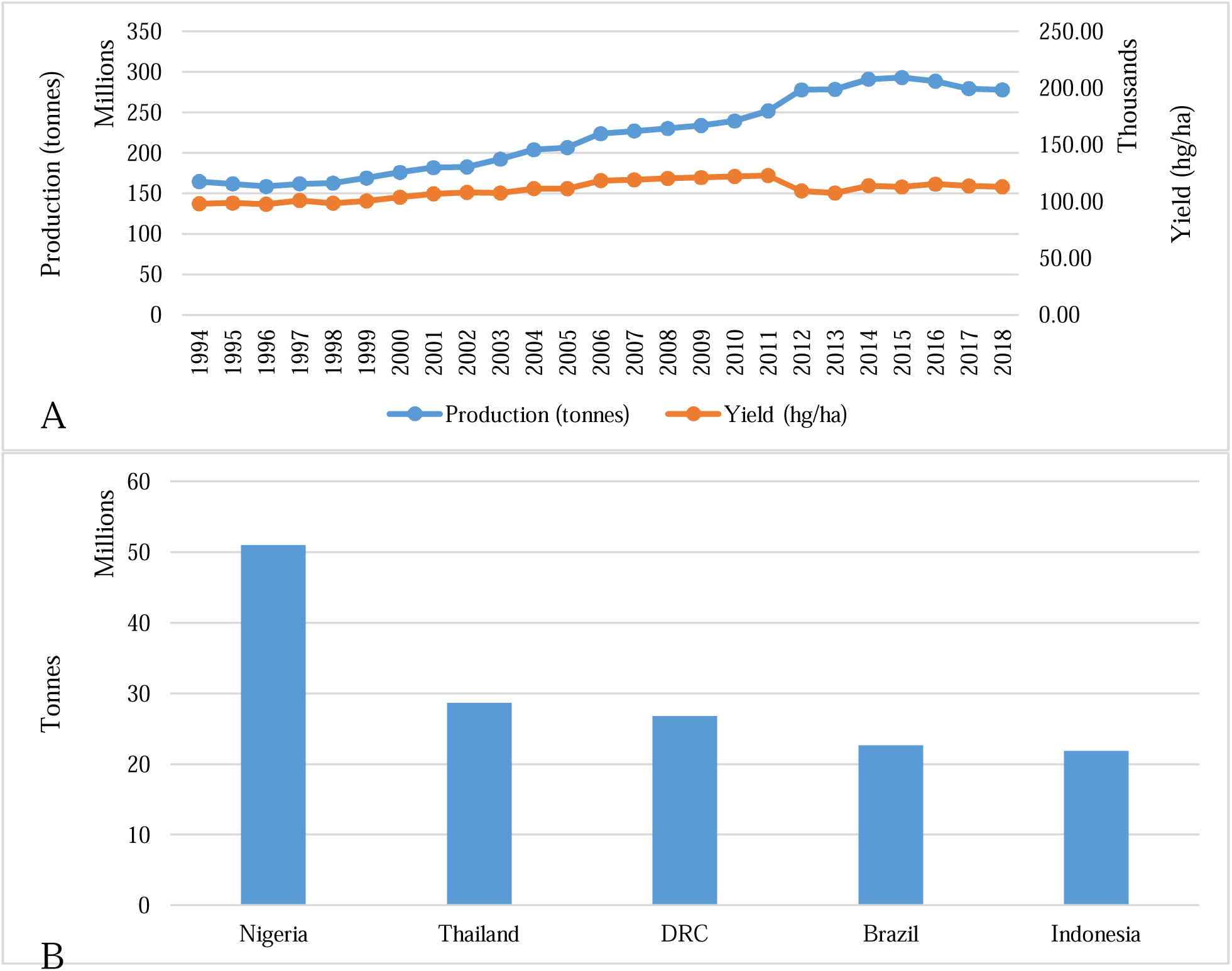
World cassava production statistics. (A) production/yield quantities of cassava in the world from 1994 – 2018. (B) Top 10 producers of cassava from 2008 – 2018. Data source: FAOSTAT [9].

Cassava root comprises of three well defined tissues, namely, periderm, cortex and parenchyma. The periderm; the outer layer of the root, sheds off as the root eventually grows and ages, and constitutes a thin layer of cells which comprises approximately 3% of the total weight of the root. The cortex is comprised of three different cells namely; cortical parenchyma, sclerenchyma and phloem cells, with these group of tissues constituting approximately 11 – 20% of the root weight. The edible portion of the root (parenchyma) constitutes an average of 85% of the total weight [3, 4]. It comprises of the xylem vessels which are radially distributed in a matrix of starch containing cells [3, 4]. Cassava comprises of a considerable amount of vitamin C (25 mg/100g), phosphorous (40 mg/100g), and calcium (50 mg/100g) [27] while the concentration of proteins, riboflavin, thiamin and niacin in cassava is very low making it the one of highest sources of carbohydrates among tuber crops [5]. The carbohydrate content of cassava ranges from 64 – 72% starch (amylose and amylopectin) which is structurally different from that found in cereal, in its branch chain length distribution, amylose content and its granular structure. Approximately 17% of sucrose is also found predominantly in the sweet varieties and small quantities of fructose and dextrose have also been reported. The protein content is determined as between 1 – 2%, with low essential amino acid profiles; particularly methionine, tryptophan and lysine, whilst conversely possessing a high dietary fiber content (3.40–3.78% soluble, and 4.92– 5.6% insoluble) [6, 7].

### Cyanogenic glycosides and Cyanide

Cyanogenic glycosides are a large group of secondary metabolites which are distributed across the plant kingdom [8]. Cyanogenic glycosides are present in all parts of the plant with the leaves having the highest concentration [9]. According to Kotopka and Smolke [10], these compounds act as chemical defenses produced by the plants as a deterrent against pathogenic organisms and the activities of herbivores. Structurally, cyanogenic glycosides comprise of a core carbon which is attached to a CN group, as well as two substituent groups denoted as R_1_ (methyl, phenyl or p-hydroxyphenyl group) and R_2_ (hydrogen, methyl or ethyl group) and attached to a monosaccharadic or disaccharidic sugar via glycosidic bonding [11].

Cassava is comprised of two cyanogenic glycosides namely lotaustralin and linamarin which release hydrogen cyanide (HCN) upon destruction of the tissues as a result of mechanical damage during harvesting, or indeed chewing action of herbivores and consumers. The presence of these glycosides, especially in the tuber has been to some extent attributed to the extreme conditions in which the crop is grown, with draught being one of the parameters investigated thus far, findings from a research monitoring cassava toxicity in mozambique showed that the levels of residual cyanide tripled during draught years in comparison to the normal years[12, 24]. The breakdown of linamarin catalyzed by an endogenous β-glucosidase (linamarase) due to the disruption of cellular integrity of a plant cell leads to the formation of a cyanohydrin and a sugar (Scheme 1). The cyanohydrin which is formed, is highly unstable under neutral conditions and undergoes further decomposition to yield an aldehyde, or a ketone and cyanide [11, 13, 14]. The enzyme hydroxynitrile lyase catalyzes the breakdown of the cyanohydrin formed into a carbonyl compound and hydrogen cyanide [15] (Scheme 1). The toxicity of a cyanogenic glycoside is as a result of its degradation catalyzed by its endogenous β-glucosidase to yield hydrogen cyanide, which would eventually lead to acute cyanide poisoning (LD_50_ of 1.52 mg/kg for oral administration) [28]. The following clinical symptoms; drop in blood pressure, dizziness, headache, mental confusion, blue colouration of skin due to lack of oxygen, twitching and convulsion, rapid pulse, stupor are usually presented in cases of acute cyanide poisoning [11].

High levels of cyanide intake associated with the chronic consumption of cyanogenic glycoside (from cassava *etc*.) are reported to lead to diseases such as iodine deficiency disorder, tropical ataxic neuropathy and konzo [16, 17].

**Scheme 1.**
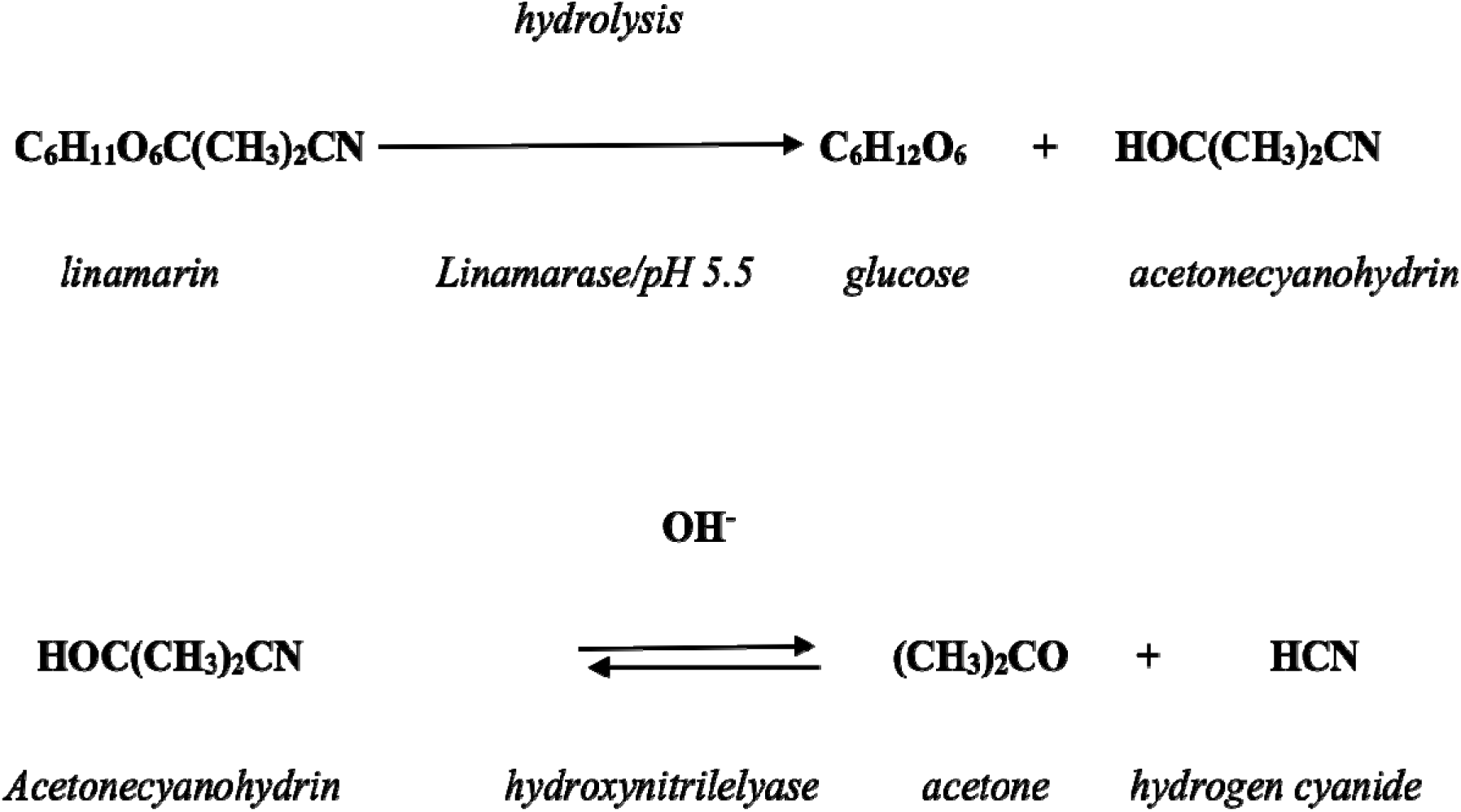
Hydrolysis of linamarin adopted from Idibie, 2006 [15]

Cassava being of a lower nutritional value than other staple foods consumed in subsaharan Africa and vitamin A deficiency being a major hindernace to improved nutrition, prompted the biofortification of cassava, giving rise to the genetically engineered pro-vitamin A cassava developed under the IITA-HarvestPlus program. This was rationalised to partially address the vitamin A deficiency affecting much of the subsaharan Africa population, with approximately 23,500 child mortalities annually in Kenya as a result of micronutrient deficiencies, with school children often suffering from sub-clinical vitamin A deficiency [17]. Herein, we determine the levels of residual hydrogen cyanide and β-carotene content as yellow flesh cassava UMUCAS 38 (TMS 01/1371) is being processed from tuber into confectionary products whilst NR 8082 is used as control sample.

## Materials and methods

### Materials

Acetone, hyflosupercel (celite), 3mm whatman filter paper, vacuum filtration equipment, UV visible spectrophotometer (Jenway 6300, Staffordshire, UK). All chemicals within this study were purchased from Sigma Aldrich (1 Friesland Drive Longmeadow Business Estate 1645 Modderfontein South Africa).

### Sample preparation

Freshly harvested roots of the two experimental cultivars UMUCASS 38 (TMS 01/1371) and NR 8082 (Control) were obtained from the Cassava Programme of National Root Crops Research Institute (NRCRI), Umudike, Nigeria. The samples were processed into high quality cassava flour (HQCF) following the methods described by Onabolu et al [18] and oven dried at a temperature of 115°C for 6 hours. The HQCF sample was furthered processed into consumer products.

### Carotenoid Determination

The extraction with acetone for carotenoid analysis developed by Rodriguez-Amaya and Kimura [19] was used for the determination of the total β-carotene content of the samples. 5 mg of the sample was ground with the aid of hyflosupercel (3.0 g) in 50ml of cold acetone and vacuum filtered. The filtrate was extracted using 40 ml petroleum ether (PE). Saturated sodium chloride was used to prevent the formation of emulsion. The lower aqueous phase was discarded while the upper phase was collected and filtered through 15g of anhydrous sodium sulfate to eliminate residual water. The seperating funnel was washed with PE and the flask was made up to 50 ml. The absorbance of the solution was measured at 450 nm and the total carotenoid content was calculated using the Beet-Lambert law (Equation 1).

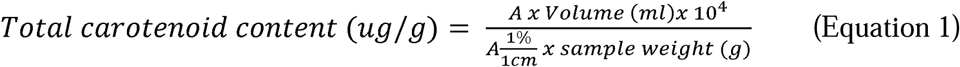

### Cyanide Determination

The simple picrate paper method was used to determine the levels of residual hydrogen cyanide [20]. 100 mg of the sample was placed in a flat-bottomed plastic bottle containing the enzyme (linamarinase), buffer and picrate paper. The contents were left to incubate in the dark for 24 hours at room tempeature. The picrate papers darkened as a result of cyanide production were then placed in test tube with 5ml of distilled water. The sample was allowed to stand at room temprature for 30 minutes. The UV absorbance was determined at a wavelength of 510 nm and total cyanide content calculated according to Equation 2.

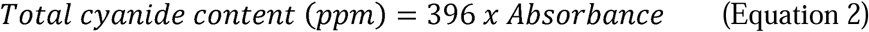

### Statistical Analysis

Paired t-tests were carried out to compare the levels of cyanide and β-carotene in the different samples using Prism 8 (Graph Pad software LLC). ANOVA was carried out using the Statistical Package for Social Sciences (SPSS), version 22. Statistical significance was set at *p<* 0.05.

## Results

### Cyanide determination

Fresh NR 8082 carried the highest cyanide concentration (44.10±0.14 ppm) while the fresh UMUCASS 38 had a value of 43.02±0.02 ppm (Figure 1). NR 8082 flour was determined as having the highest cyanide level (18.01±0.01 ppm) while the UMUCASS 38 had the least (17.02±0.02 ppm). The cookie sample showed the highest concentration (10.00±0.00 ppm) as compared to the cake sample (7.10±0.14 ppm). In addition, the NR 8082 variety had significantly higher cyanide concentration (*p*<0.05) than the yellow flesh (Fig. 2).

**Fig. 2.**
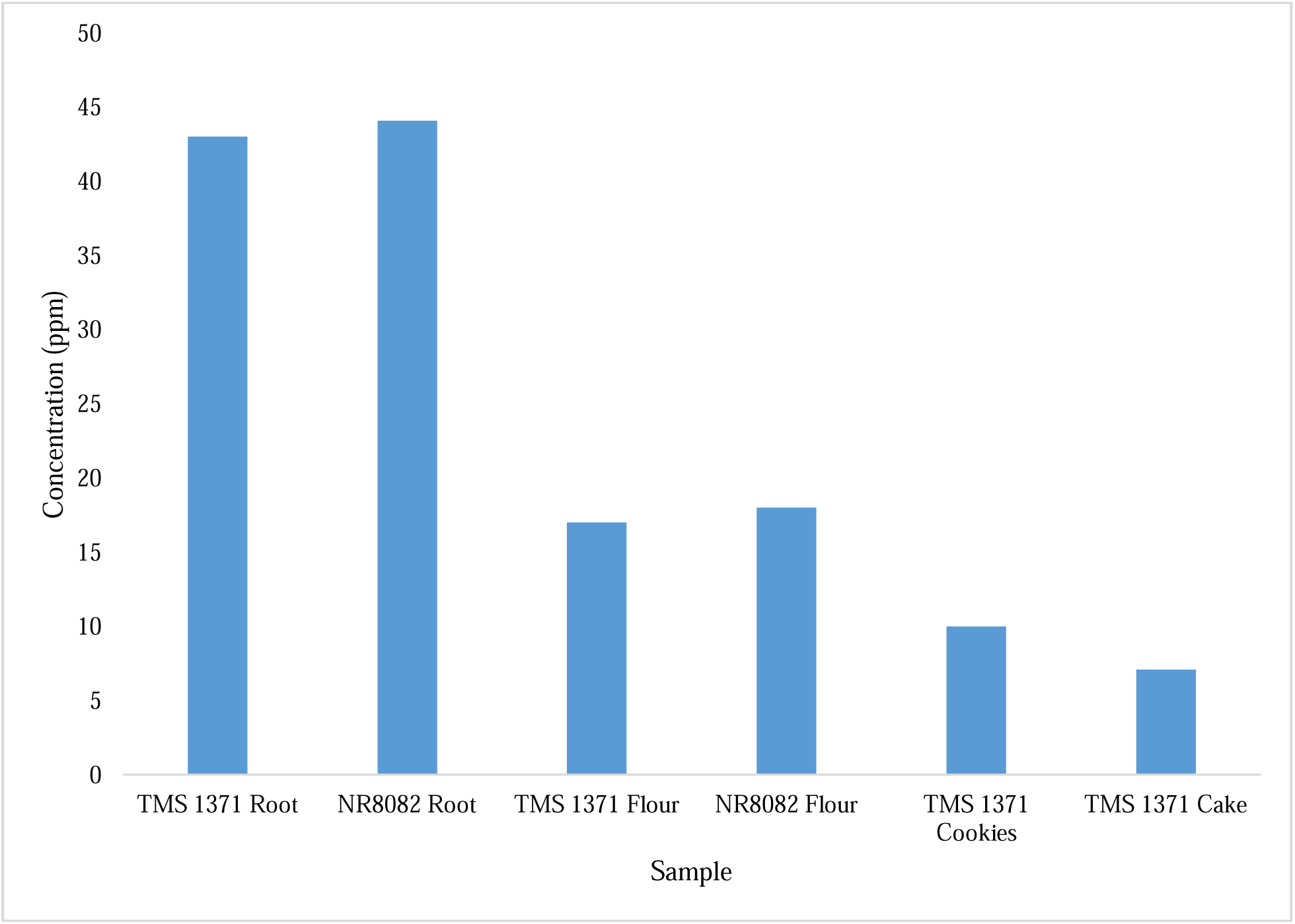
Levels of residual cyanide in roots and products determined using simple picrate paper method.

### Carotenoid determination

Fresh UMUCASS 38 possessed a carotenoid content of 6.53±0.02 µg/g compared to that of the NR 8082 variety (1.17±0.02 µg/g). The products retained a portion of the β-carotenoids after production; the cake sample had a residual β-carotene concentration of 2.84±0.04 µg/g whilst that of the cookie sample was determined at 2.15±0.01 µg/g (*p*<0.05) (Fig. 3).

**Fig. 3.**
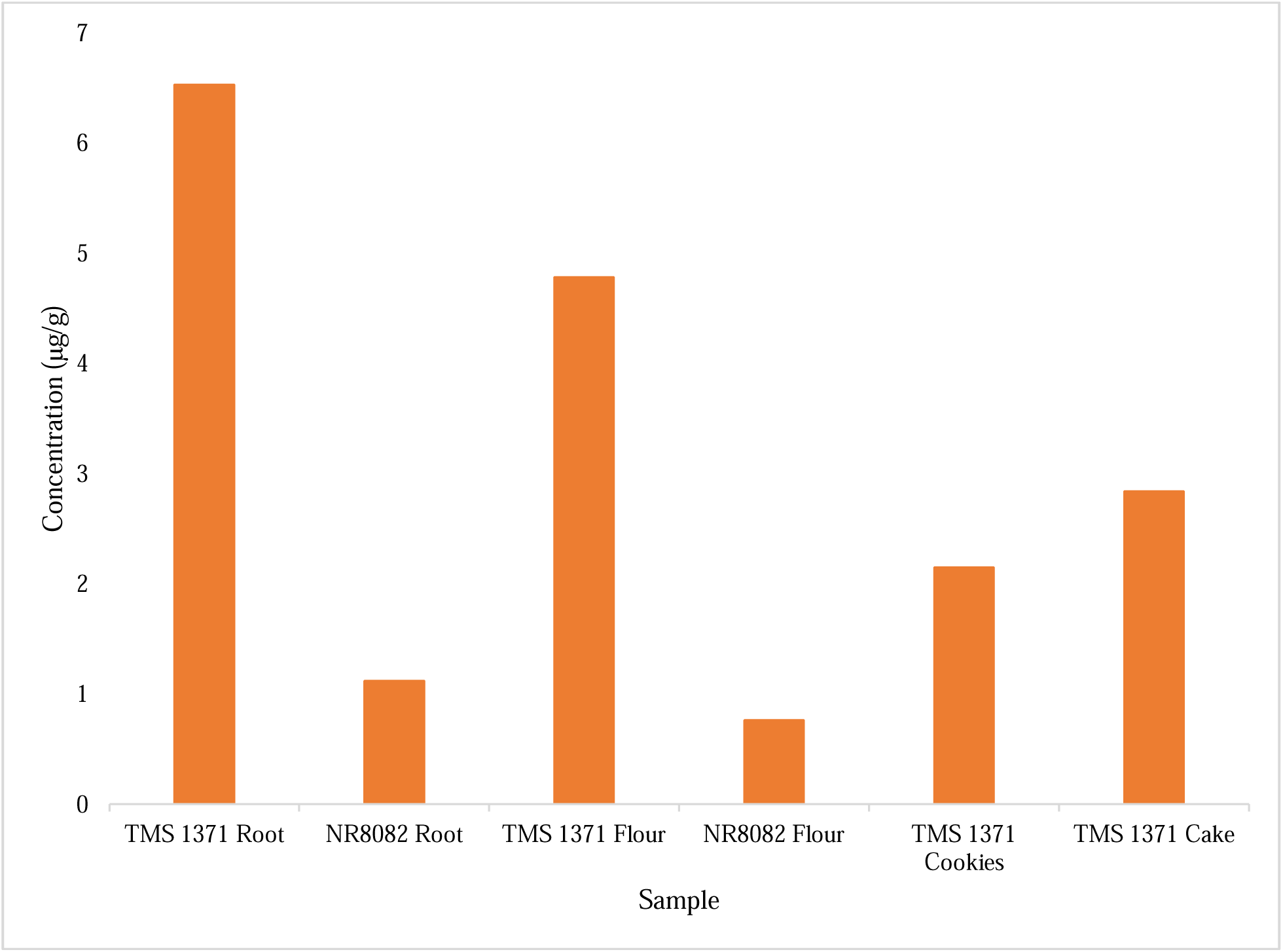
Levels of *β*-Carotene in roots and products determined using the extraction with acetone method for carotenoid analysis developed by Rodriguez-Amaya and Kimura.

## Discussion

Chronic exposure to cyanide causes a myriad of cardiac, neurological and metabolic dysfunctions which can be fatal [21]. As a result of concern regarding the levels of potential residual cyanide remaining in cassava after processing, the roots were classified according to their potential toxicity to humans and animals as non-toxic (less than 50mg HCN kg^−1^ in fresh root), moderately toxic (50-100mg HCN kg^−1^ in fresh root) and highly toxic (above 100mg HCN kg^−1^ in fresh root) [22]. The lethal dose of cyanide in humans is in the range of 0.5 to 3.5 mg/kg body weight [23, 24]. The level of cyanide in the flour within this study was reduced by almost 60% as a result of the method of food processing. The products possessed lower levels of cyanide; acceptable according to the WHO standard of 10 ppm [13]. This standard was reached as a result of the lack of quantitative and epidemiological information to estimate a safe level. However, the JECFA committee concluded that upto a level of 10 mg HCN/kg body weight (10 ppm) in the codex standard of cassava flour is not associated with acute toxicity [25]. The low cyanide levels in the products was as a result of the processing method which involved the peeling, grating and subsequent oven drying to produce HQCF. The low cyanide levels in the products suggest that the food products may not be highly toxic to consumers when employing the WHO standard as a benchmark [13]. The body has several pathways for the detoxification of cyanide, and this primarily involves the conversion of soluble thiocyanate (SCN^−^) by the enzyme rhodanase [25]. Lesser pathways of metabolism include the complexation of cyanide with cobalt in hydroxocobalamin to form cyanocobalamin (Vitamin B12) [25].

The consumption of these cassava varieties as a staple food must be complemented by a diet rich in protein from exogenous sources due to the low protein content of cassava itself; the findings of the current study showed a reduction in cyanide and β-carotene levels in the processed products (Cookies, 10.00±0.00 ppm; Cake, 7.10±0.14 ppm) and (Cookies, 2.15±0.01 µg/g; Cake, 2.84±0.04 µg/g) respectively (Figs. 2 & 3). The levels of β-carotene after processing using a method which has been confirmed to reduce cyanide levels at the expense of leaching or destruction of essential nutrients such as vitamin C, β-carotene (vitamin A precursor) and vitamins B (riboflavin, niaci and thiamine) suggests that the consumption of yellow root cassava UMUCAS 38 does indeed contribute to the recommended daily allowance of vitamin A [24]. The continuous consumption of cassava-based products without sufficient protein intake would limit protein synthesis, thus leading to stunted growth in children [22].

Carotenoids, the colourful plant pigments, some of which the body can convert to vitamin A, are also powerful antioxidants that have been suggested to contribute to the resistance against certain forms of cancer and heart diseases, and also enhance immune response to infections [22]. The yellow cassava species investigated had significantly higher carotenoid quantities than the white variety, thus this may confer antioxidant potential [22]. The predominant carotenoid in yellow cassava being β-carotene, suggests a need for dietary supplementation as the consumption of this yellow root cassava may not meet the recommended daily allowance (RDA) for vitamin A in men (750 – 900 µg daily), women (700µg daily) and children (400 – 600 µg daily) [22, 26].

Fresh UMUCASS 38 had the highest carotenoid content (6.53 µg/g) while the NR 8082 variety (1.12 µg/g) had relatively low carotenoid content in comparison. There was a decrease in the carotenoid content in the flour level as a result of exposure to light and heat treatment. The products retained a portion of the β-carotenoids after heat treatment, this could be as a result of the ingridients used in the making of the product which includes eggs, a known source of vitamin A. There was also a significant decrease in the HCN levels which can be attributed to further heat treatment i.e. baking, mixing of the sample which could have lead to the release of the enzyme linamarinase.

## Conclusion

This study determined the levels of residual hydrogen cyanide and β-carotene as the yellow flesh cassava UMUCAS 38 (TMS 01/1371) is processed from tuber into confectionary products. The results obtained from the study showed that the processed yellow root variety had low levels of residual cyanide. The UMUCASS 38 variety retained relatively significant quantities of β-carotene after (Cookies, 2.15±0.01 µg/g; Cake, 2.84±0.04 µg/g) processing through peeling, grating, heat treatment (oven drying), milling, these processes are known to diminish nutritional value as well as cyanide content. The consumption of the pro vitamin A cassava variety should be encouraged as the findings herein demonstrate the viable food safety of the cassava-based products for human consumption as well as the need to supplement vitamin A from exogenous sources to combat cases of vitamin A deficiency in regions where cassava is a staple food. Based on the findings from this study we suggest that more research should be carried to further improve the β-carotene content of these biofortified cassava varieties.

## Limitations

- Using the simple picrate paper method for cyanide determination is limited by the rate of reaction of approximately 16 – 24 hrs for completion, the chemicals require special handling and storage, the results obtained can sometimes be indefinite. Hence, the dissolving of the chromophore from the picrate paper for a quantitative determination using a spectrophotometer.

## Abbreviations

FAO: Food and Agriculture Organization
IITA: International Institute of Tropical Agriculture
HQCF: High Quality Cassava Flour
ANOVA: Analysis of Variance
SPSS: Statistical Package for Social Sciences
DRC: Democratic Republic of Congo
P.E.: Petroleum ether
JECFA: Joint FAO/WHO Expert Committee on Food Additives.

## Ethics approval and consent to participate

Not applicable

## Availability of data and material

All data generated or analysed during this study are included in this published article.

## Funding

Not applicable.

### Acknowledgements

CSO and PNO thanks the late emeritus Professor Howard Bradbury for providing the cyanide determination equipment.

## Consent for publication

All authors consent to the publication of this manuscript.

## Author’s contributions

Chiemela S. Odoemelam (CSO) and Polycarp N. Okafor (PNO) carried out the analyses. PNO, Dawn Scholey (DS), Emily Burton (EB) and Philippe B. Wilson (PBW) contributed to experimental design. Benita Percival (BP), CSO and PBW carried out the statistical analyses. CSO developed the first draft of the manuscript. Zeeshan Ahmad (ZA), Ming-Wei Chang (MWC), EB, DS, PNO and PBW contributed to supervision, manuscript development, analysis of results and revision of manuscript drafts.

